## Supplemental information for "Erythrocyte CD55 facilitates the internalization of *Plasmodium falciparum* parasites"

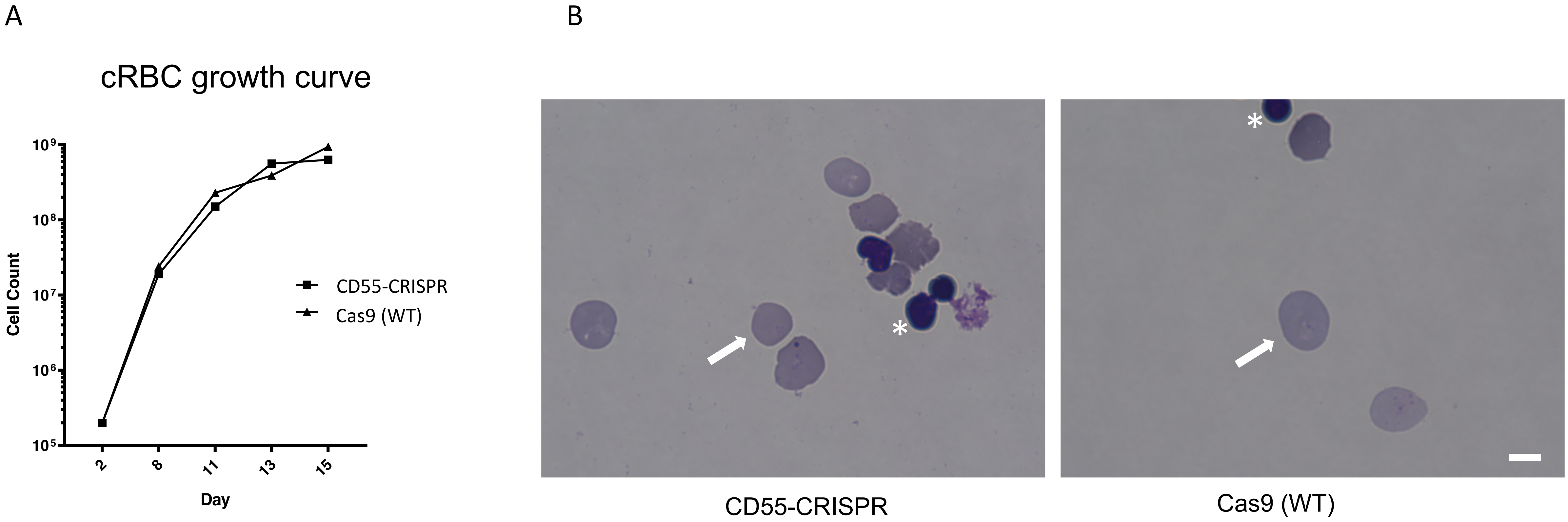


**Figure S1: Deletion of *CD55* does not impact growth or maturation of cRBCs derived from CD34+ primary human HSPCs.** (A) Growth rate of *CD55*-null erythroid progenitors from CD34+ HSPCs over 15 days of differentiation, compared to isogenic wild-type cells. Results are from one representative experiment. (B) Day 20 cRBCs stained with May-Grünwald and Giemsa, and visualized by light microscopy. Enucleation rate of ~90.0% was observed in both *CD55*-null and Cas9 (WT) cRBCs. White arrows indicate enucleated cells. Asterisks indicate free nuclei. Scale bar 5 µM.

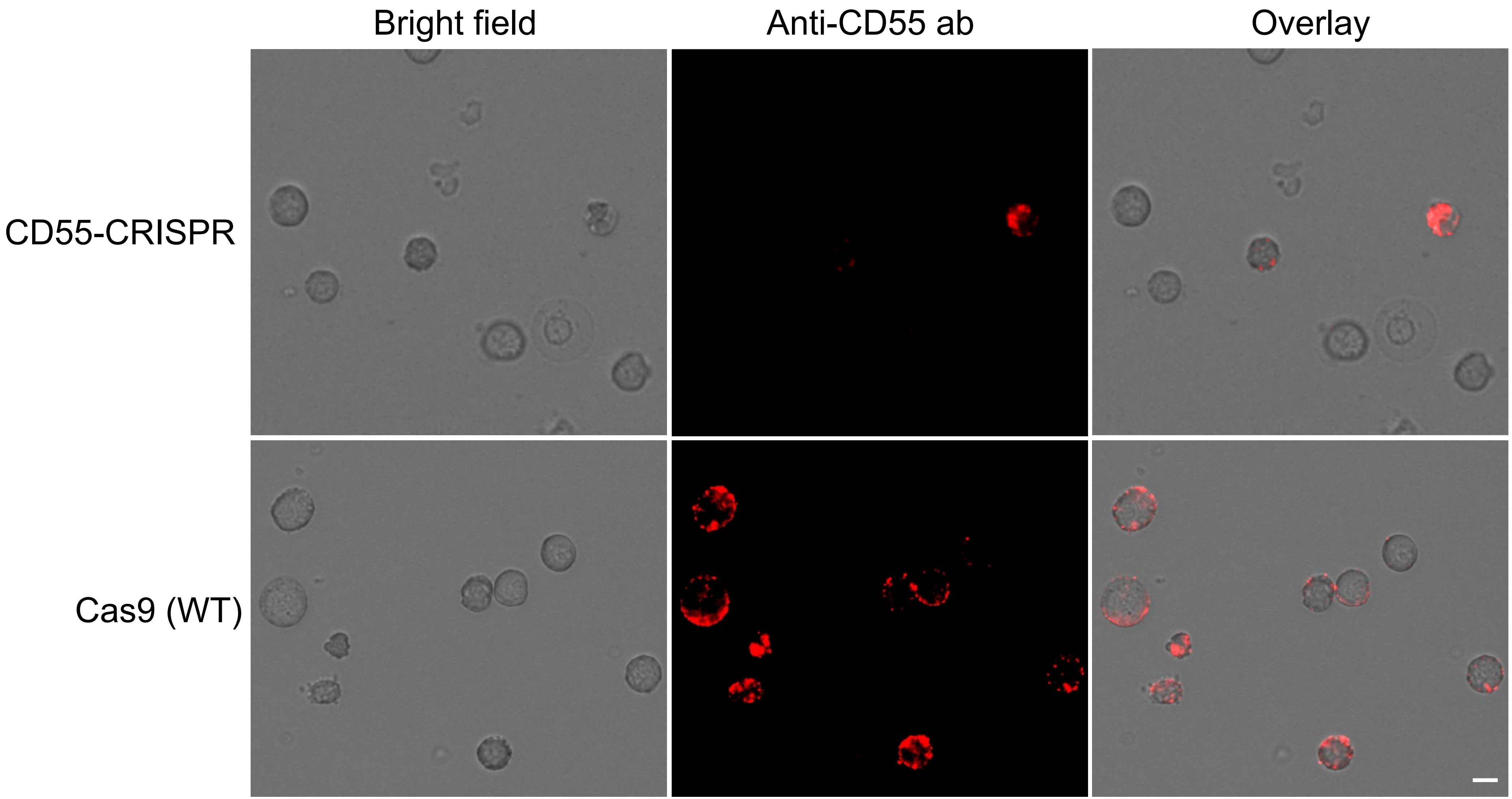


**Figure S2: Expression of CD55 on cRBCs.** Immunofluorescence assays showing the expression of CD55 on the surface of CD55-CRISPR and Cas9 (WT) cRBCs using anti-CD55-PE antibody. Scale bar 5 μM.

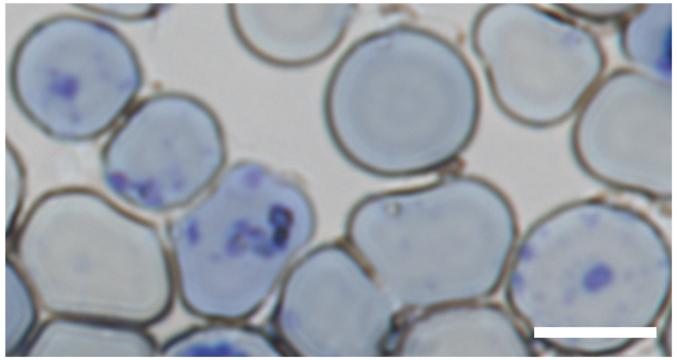

**Figure S3: Reticulocyte staining of day 17 cRBCs.** Reticulocytes are indicated by the purple reticular staining pattern. Scale bar 5 µM.

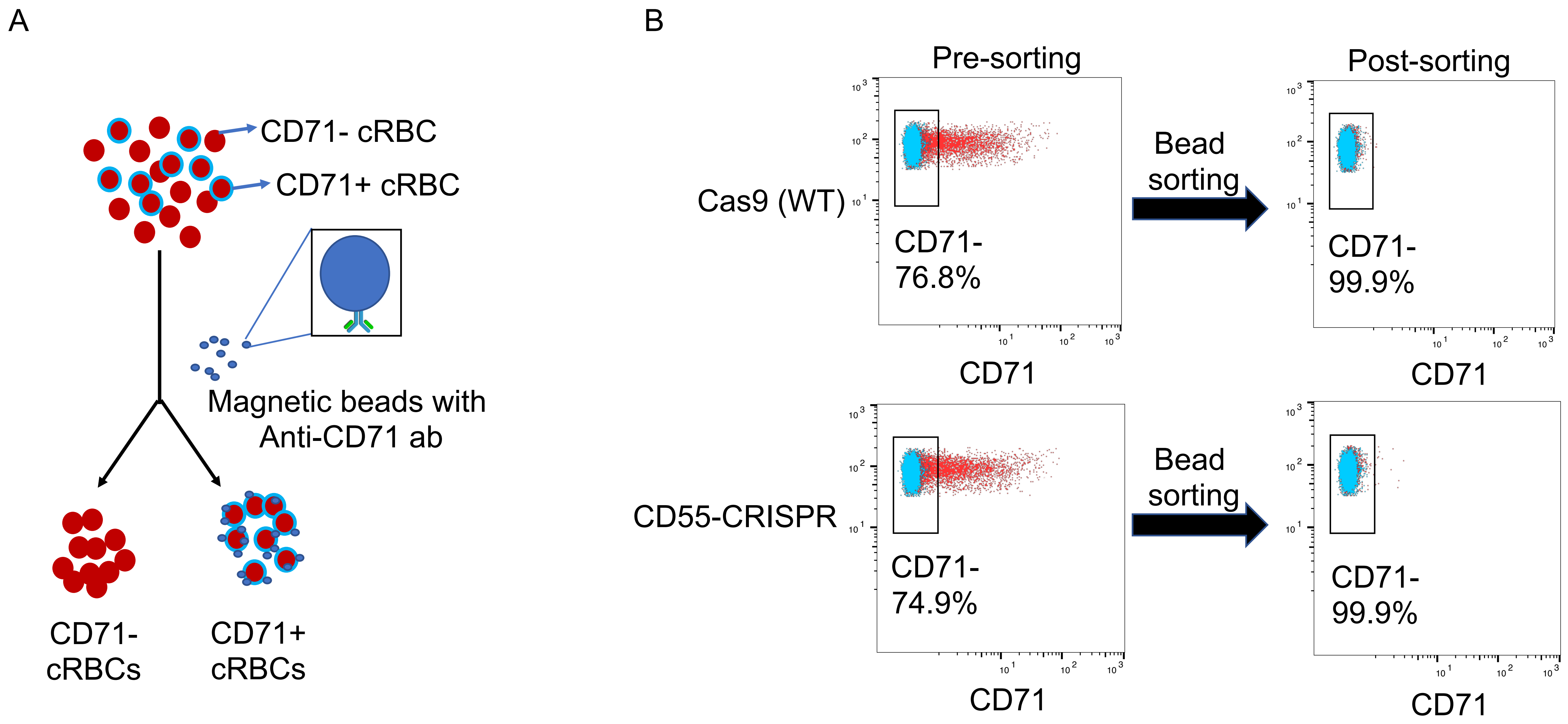


**Figure S4: Enrichment of CD71- cRBCs by immunolabeled magnetic beads.** (A) Schematic showing the separation of CD71+ cRBCs from CD71- cRBCs using anti-CD71 antibody-immobilized magnetic beads. (B) Flow cytometric analysis of CD71 expression on cRBCs pre- and post- sorting using anti-CD71 antibody immobilized magnetic beads

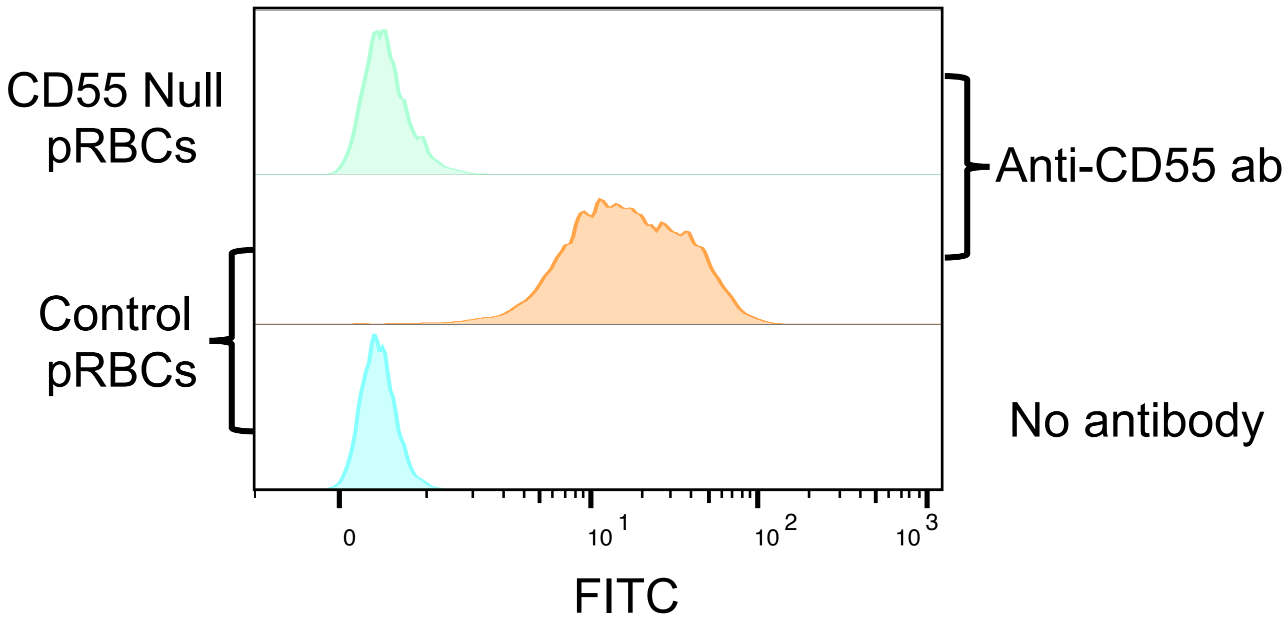


**Figure S5: Specificity of rabbit polyclonal anti-CD55 antibody.** Flow cytometric analysis of control and CD55-null RBCs incubated with rabbit polyclonal anti-CD55 antibody. The anti-CD55 polyclonal antibody detects CD55 on wild-type pRBCs but not CD55-null RBCs.

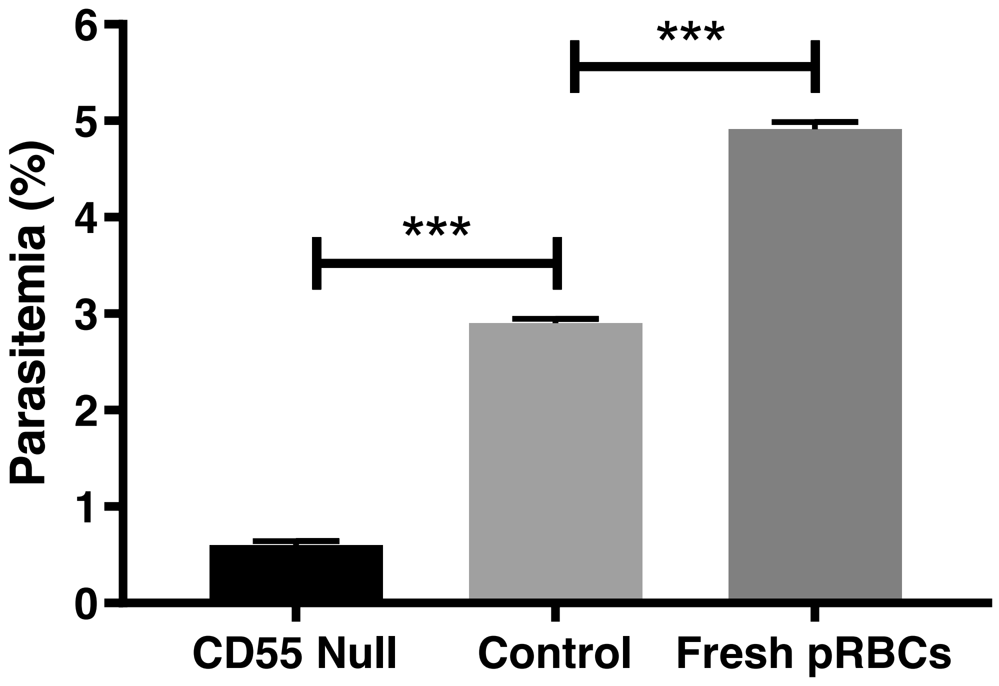


**Figure S6: CD55-null pRBCs are refractory to invasion by *P. falciparum***. Invasion efficiency of *P. falciparum* strain 3D7 in previously cryopreserved CD55-null pRBCs from rare Inab donor, control wild-type pRBCs that had been similarly cryopreserved, or fresh pRBCs. n=3 technical replicates, error bars indicate SD; ***, $p<0.05$.
